# Etomidate as an Effective Alternative Method of Euthanizing Adult Zebrafish (*Danio Rerio*)

**DOI:** 10.1101/2024.11.02.621650

**Authors:** K Schwartz, K Byrd, M Huss, K Jampachaisri, P Sharp, C Pacharinsak

**Author notes:** Corresponding author: Kenzie Schwartz. **Author Contribution Statement Kenzie Schwartz:** Investigation, Writing-Original Draft, Methodology. **Kyna Byrd:** Investigation. **Monika Huss:** Writing-Review and Editing, Investigation. **Katechan Jampachaisr**i: Formal analysis. Patrick Sharp: Writing-Review and Editing, Investigation. **Cholawat Pacharinsak:** Writing-Review and Editing, Investigation, Methodology.

## Abstract

We assessed the efficacy of etomidate in euthanizing adult zebrafish. The aims of this study were to investigate: 1) the efficacy of 2 concentrations and 2) the effects at 2 densities. Aim 1-Fish (n=10 fish/group) were immersed in either 6 mg/L (Eto-6) or 10 mg/L (Eto-10) etomidate. Aim 2– either 5 or 10 fish were immersed in Eto-6. Tanks were video recorded and scored by two blinded observers. The parameters monitored included: 1) loss of righting reflex (LORR); 2) undulation cessation; 3) cessation of operculation (COO); and 4) loss of startle reflex. After the 30 minute immersion, fish were transferred to recovery tanks and monitored once every 5 minutes for 60 minutes. Results – Aim 1, the time to undulation cessation, COO, and loss of startle reflex was significantly shorter in the Eto-10 group and there was no difference in LORR. Aim 2, the time to LORR and undulation cessation was significantly shorter in the 5 fish/L group and there was no significant difference in time to COO or loss of startle reflex. No aversive behaviors were noted and fish in all groups were effectively euthanized by 30 minutes. This study indicates that Eto-6 or Eto-10 effectively euthanizes adult zebrafish.

## Introduction

Zebrafish are one of the most commonly used research animals, with an estimated 3,250 institutes in 100 countries using over 5 million zebrafish annually [1]. These numbers are expected to grow with gene editing technology advances such as CRISPR/cas9. With this vast number of fish euthanized daily all over the world there remains a significant need to improve the standard euthanasia methods. For euthanasia to be deemed humane, certain conditions must be met. These include ensuring minimal pain and distress for the animal, inducing rapid unconsciousness with minimal restraint leading to irreversible death, and ensuring reliable central nervous system depression. Moreover, the process should be reproducible, simple, safe, and avoid emotional strain for the operator. It should also be economically viable and tailored to factors such as species, strain, age, number, and health of the animal. In research settings, it should not compromise post-mortem analysis [2,3]. The current standard for euthanizing zebrafish over 15 days old remains tricaine methanesulfonate (MS-222) anesthetic overdose, which can be aversive to the fish, is hazardous to human health, and thus requires special handling and preparation. Rapid chilling is another acceptable euthanasia method in the United States, while rapid chilling in European nations remain questionably humane [1,2]. Additionally, electroshock is a widely accepted euthanasia method in aquaculture, but is currently not logistically viable in laboratory animal medicine.

Currently many other anesthetic agents are being evaluated for potential use in zebrafish euthanasia by overdose. One paper found etomidate to be one of two agents, along with TBE, to be the least aversive when using hydrochloric acid as a control when comparing it to other anesthetic overdose options such as benzocaine/lidocaine HCL, 2,2,2 tribromethanol (TBE), and clove oil [4]. Lidocaine HCL and benzocaine are acidic compounds, which potentiates their ability to be aversive to fish without a buffering compound similarly to MS-222. Clove oil and one of its constituents, methyl eugenol, is a suspected carcinogen and the FDA has given guidance on not using it in aquatic species, especially if it has potential to end up in the food system [5]. We chose not to evaluate TBE due to reports of difficulty with storage and preparation [6,7]. Etomidate was chosen as our anesthetic overdose euthanasia agent due to reports of minimal aversion, ease of use, wide human safety margins, and availability.

Etomidate is a non-barbiturate hypnotic compound commonly used as an induction agent in mammalian anesthesia. Etomidate acts as an allosteric agonist on the γ-aminobutyric acid inhibitory neurotransmitter [8]. Etomidate’s binding site is located in the transmembrane site between the beta and alpha GABAA subunits. This specific binding site is associated with a Cl-ionopore and works by increasing the duration of time the Cl-ionopore is open and therefore increases GABA’s effect. An advantage for its use in mammalian anesthesia is its minimal cardiovascular and respiratory system effects [9]. It is quick acting but does not provide any analgesia [10]. Etomidate is primarily metabolized by the liver and has no active metabolites. Etomidate can also be easily stored and is readily available due to common use in human medicine.

In this study we compared 1) etomidate’s efficacy at two commonly used concentrations (6 mg/L and 10 mg/L) and 2) etomidate’s efficacy at a single concentration (6 mg/L) used to euthanize two separate fish densities (5 fish/L or 10 fish/L). We hypothesized whether Eto-10 would result in a shorter time to LORR, undulation cessation, COO, and loss of startle reflex than Eto-6. We also hypothesized fish euthanized in Eto-6 at a density of 5 fish/L would have shorter times to LORR, undulation cessation, COO, and loss of startle reflex than 10 fish/L in Eto-6.

## Materials and Methods

### Ethics statement

This study was designed with the 3Rs in mind to test the efficacy and assess etomidate’s aversion using different concentrations and fish densities as a euthanasia method. All procedures were approved by Stanford University’s IACUC, an AAALAC-accredited institution.

### Animals and husbandry

Thirty mixed sex wild-type AB zebrafish from a single batch of embryos (Sinhuber Aquatic Research Laboratory, Corvallis, OR) were used for this study. Zebrafish racks in this facility undergo regular PCR screening for infectious agents (*Aeromonas hydrophila, Edwardsiella ictaluri, Flavobacterium columnare, Mycobacterium* spp., *Ichthyophthirius multifiliis, Pleistophora hyphessobryconis, Pseudocapillaria tomentosa, Piscinodinium pillulare, Pseudoloma neurophilia*, and *Saprolegnia brachydanis*) and no potentially pathogenic organisms were detected on our rack at the time of this experiment. All fish used appeared healthy and behaving appropriately.

Zebrafish were group housed based on the Stanford IACUC guidelines at a stocking density of 5 to 10 fish/L on a single recirculating aquaculture system rack (Aquaneering, San Diego CA) in a housing room with a 14:10 h-light:dark cycle. The system rack was fitted with a heater, 50-um prefilter, 25-um mechanical filter, fluidized bed biologic filter, carbon filter, and UV lamp providing a minimum of 100,000 mW/s/cm2. A daily 10% water volume change was performed daily with calcite-filtered reverse osmosis water. Water temperature and chemistry were maintained within standard ranges established as species-appropriate (25 to 29 °C, pH 6.5 to 8.5, dissolved oxygen > 6 mg/L, conductivity 500 to 2,000 uS, ammonia < 0.02 ppm, nitrite < 0.5 ppm, and nitrate < 50 ppm). Fish were fed a diet of hatched in house *Artemia* (E-Z Egg, Brine Shrimp Direct, Odgen, UT) and commercially available powdered feed (GemmaMicro 300, Skretting USA, Tooele, UT) twice daily.

### Experimental design

Each fish was netted and placed in a 1-L clear, polycarbonate tank (Tecniplast USA, Inc., West Chester, PA) containing the original tank system water. The pH of each etomidate euthanasia preparation was tested in a pilot study and was within the tolerable range for zebrafish (7.0-7.3).

**Aim 1**. Zebrafish were randomly netted from their home tanks and randomly assigned to 1 of 2 treatment groups: Eto-6 or Eto-10 immersed at a density of 10 fish/L.

**Aim 2**. Zebrafish were randomly netted from their home tanks and randomly assigned to 1 of 2 treatment groups: 5 fish/L or 10 fish/L in Eto-6.

Fish were only placed in tanks with other fish from their same home tanks. After transferring the fish to the tank by net, etomidate was added through a syringe and disbursed as evenly as possible to obtain the desired concentration. Video was used to record the tanks for later analysis by blinded individuals experienced with fish euthanasia. A table tap was performed roughly every 5 minutes to access the startle reflex. After 30 minutes of immersion, each fish was removed from the tank and placed in a fresh system-water tank for an additional 60 minutes. Fish were monitored for signs of life (righting, operculation, body movement, and response to tap) every 5 minutes. Successful euthanasia was defined as reaching irreversible stage IV anesthesia for at least 60 min after transfer from the euthanasia tank to fresh system-water tank.

### Data Criteria

Fish video footage was observed by two blinded observers experienced with zebrafish euthanasia. Time to loss of righting reflex (LORR), undulation cessation, cessation of operculation (COO), and loss of startle reflex were recorded by each observer. LORR was defined as the inability to remain upright. Undulation cessation was defined as complete loss of purposeful movement forward. COO was defined as loss of opercular movement or buccal pumping. Loss of startle reflex was defined as no reaction or movement to the table being tapped every 5 minutes. Times were then averaged between the two observers and compared between groups. Displays of aversive behaviors (erratic movement, hyperlocomotion, piping at the surface, twitching) were also noted if seen.

### Statistical analysis

Times until LORR, undulation cessation, COO, and loss of righting reflex were analyzed by using the Mann–Whitney *U* test. Data are expressed as median ± interquartile range (IQR). A *P* value less than 0.05 was considered significant. Analysis was performed by using R Core Development Team, 2015.

## Results

### Aim 1

The median times to LORR, undulation cessation, COO, and loss of startle reflex are summarized along with the IQR in a box and whisker graph in Figure 1. There was no significant difference in time to LORR between the Eto-6 and Eto-10 groups, with a median of 24.0 (IQR, 21.3 to 26.5) seconds (sec) in the Eto-6 group and 24.0 (IQR, 23.0 to 26.5) sec in the Eto-10 group. The time to undulation cessation was significantly shorter in the Eto-10 group with a median time of 30.0 (IQR, 27.0 to 32.3) sec compared to the Eto-6 group with a median time of 60.5 (IQR, 45.5 to 62.0) sec (P= 0.028). The time to COO was also significantly shorter in the Eto-10 group with a median of 99.0 (IQR, 92.5 to 101.8) sec compared to the Eto-6 group with a median of 112.0 (IQR, 106.2 to 114.0) sec (P= 0.005). The time to loss of startle reflex was also significantly shorter in the Eto-10 group, with a median of 360.0 (IQR, 36.0 to 465.0) sec compared to the Eto-6 group with a median time of 720 (IQR, 480 to 720) sec (P=0.038).

**Figure 1.**
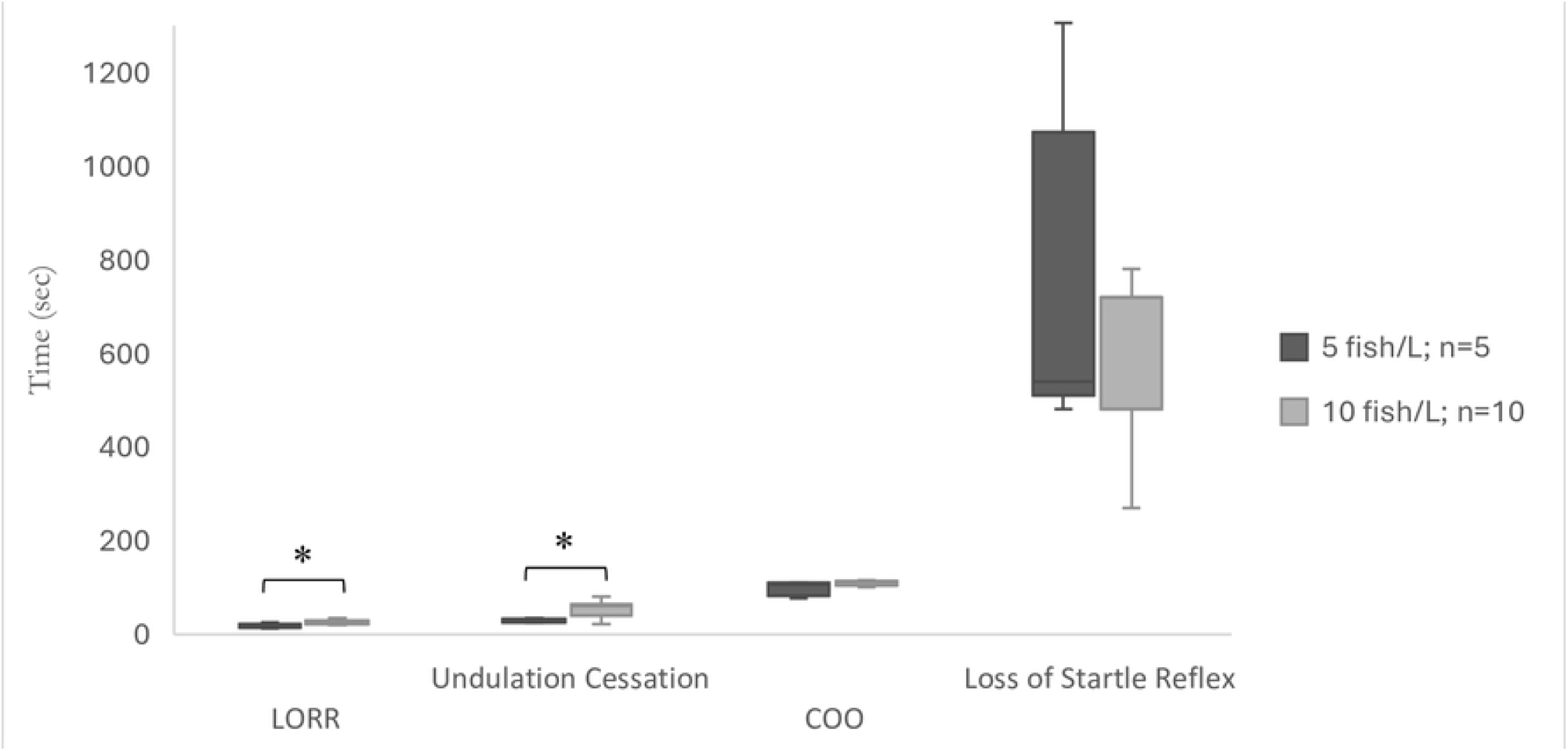
Aim 1-time to LORR, undulation cessation, COO, and loss of startle reflex (sec) in adult zebrafish after immersion in Eto-6 or Eto-10. Medians with interquartile ranges are indicated. Statistical significance was assessed by Mann-Whitney U test. * value is significantly different from the other group (P < 0.05).

### Aim 2

The median times to LORR, undulation cessation, COO, and loss of startle reflex are summarized along with the IQR in a box and whisker graph in Figure 2. The time to LORR was significantly shorter in the 5 fish/L group with a median time of 17.0 (IQR, 15.0 to 19.0) sec compared to the 10 fish/L group with a median time of 24 (IQR, 23.3 to 26.5) sec (P=0.02). The time to undulation cessation was also significantly shorter in the 5 fish/L group with a median time of 30.0 (IQR, 25.0 to 32.0) sec compared to the 10 fish/L group with a median time of 60.5 (IQR, 45.5 to 62.0) sec (P=0.037). There was no significant difference in time to COO in the 5 fish/L group compared with 10 fish/L group with a median time of 106.0 (IQR, 89.0 to 109.0) sec in the 5 fish/L group and 112.0 (IQR, 106.2 to 114.0) sec in the 10 fish/L group. Additionally, there was no significant difference in time to loss of startle reflex in the 5 fish/L group with a median time of 540.0 (IQR, 540.0 to 840.0) compared to the 10 fish/L group with a median time of 720.0 (IQR, 480.0 to 720.0) sec.

**Figure 2.**
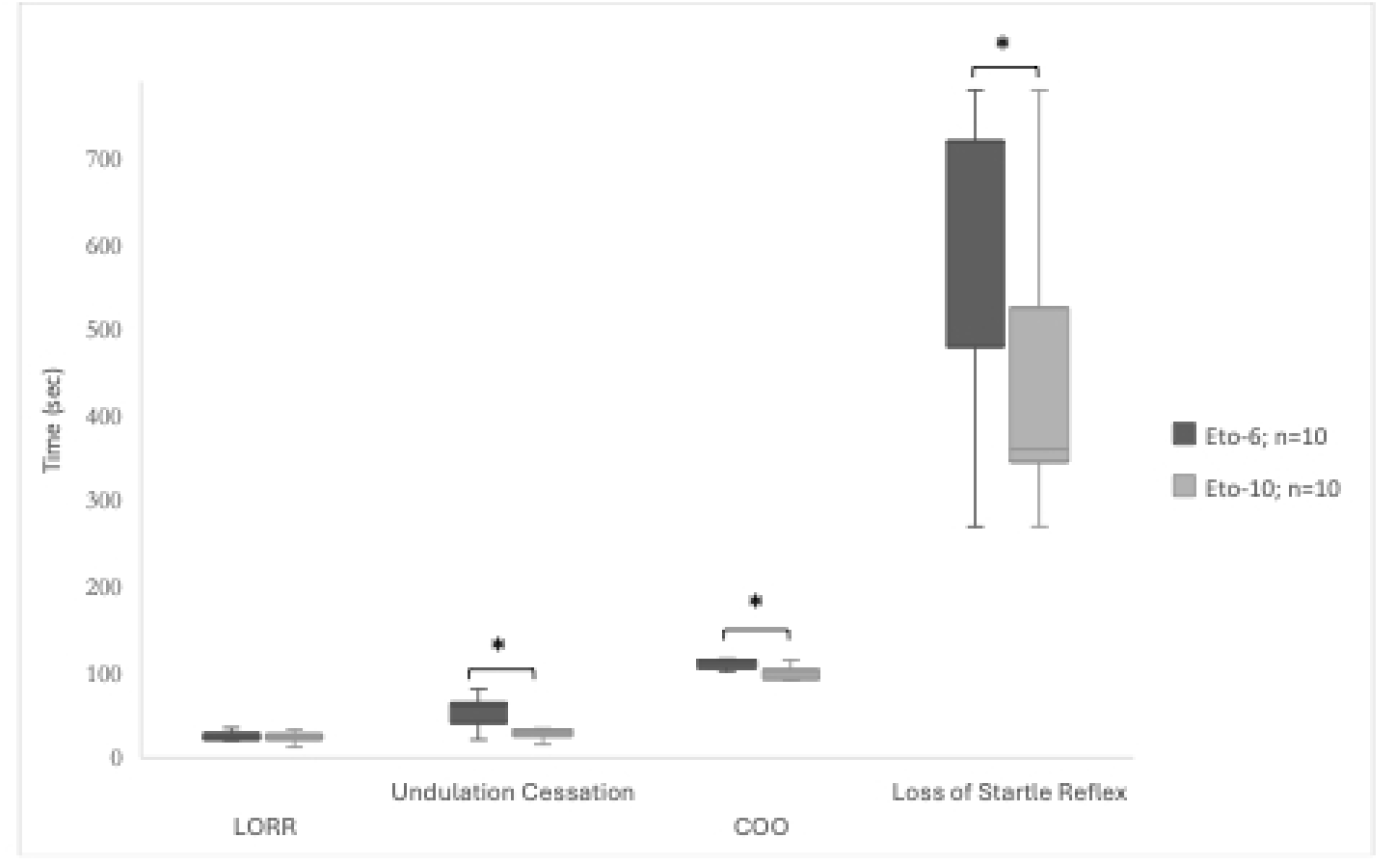
Aim 2-time to LORR, undulation cessation, COO, and loss of startle reflex (sec) in adult zebrafish after immersion in Eto-6 at densities of 5 fish/L and 10 fish/L. Medians with interquartile ranges are indicated. Statistical significance was assessed by Mann-Whitney U test. * value is significantly different from the other group (P < 0.05).

## Discussion

Results of this study indicate that for aim 1) Eto-10 and Eto-6 effectively euthanized adult zebrafish with Eto-10 resulting in shorter times to undulation cessation, COO, and loss of startle reflex. For aim 2) at 30 minutes both 5 fish/L and 10 fish/L resulted in effective euthanasia of adult zebrafish, with 5 fish/L resulting in shorter times to LORR and undulation cessation. At 30 minutes both concentrations and immersion densities effectively euthanized adult zebrafish with no signs of aversion.

For Aim 1, we found the higher concentration, Eto-10 resulted in shorter times to undulation cessation (median times; 30 sec *vs* 60.5 sec), COO (median times; 99 sec *vs* 112 sec), and loss of startle reflex (median times; 360 sec *vs* 720 sec) than Eto-6, respectively. The startle reflex was lost at a much longer time than any other parameter measured across all treatment groups. Ferreira et al also evaluated etomidate as method of euthanasia and found that Eto-6 led to a quick LORR. However, this group did not monitor for loss of startle reflex, rather loss of response to painful stimuli, which they found occurred at about 15 min, later than the other euthanasia methods being evaluated (propofol/lidocaine, clove oil and rapid chilling) [10]. Another study that evaluated etomidate at a much higher concentration (50 mg/L) saw shorter times to LORR and COO than either of the concentrations we evaluated along with a prolonged startle reflex that lasted beyond the 600 sec in which they evaluated the fish. They also noted some fish would lose the startle reflex but regain it at a later time-point, similarly to what we observed [11]. The only times recorded in our study were the final times the startle reflex was lost. The prolonged startle reflex, along with intermittent body twitching that was observed after about 20 minutes in all etomidate euthanasia groups, may result from glutamate accumulation. Myoclonic or involuntary muscle movements are a common side effect of etomidate in mammalian patients, and although the mechanism for these movements remains unclear, there has been recent evidence in Sprague-Dawley rats that it is associated with neocortical glutamate accumulation [12]. There were no differences between any of the groups in the level of twitching observed and all twitching had stopped by 30 min. The question then remains whether the fish are fundamentally brain dead at that time point and experiencing post-mortem muscle contractions. Future work utilizing an EEG would be beneficial to assess this question. Additionally, public perception of twitching should be considered as lay person may view this as conscious perception or pain in contrast to unconscious reflexive movement. The prolonged twitching and reflexive movements emphasize the importance of an adjunct euthanasia method to confirm proper brain death.

For Aim 2, animal density was assessed as a potential factor impacting the anesthetic agent’s euthanasia efficacy. We hypothesized, based on clinical experience, the agent would become less efficacious at a higher fish density (10 fish/L). We also took into consideration zebrafish housing density guidelines based on our facility’s IACUC guidelines. Euthanizing zebrafish at higher densities than their current housing density may cause unwanted stress. The Guide recommends housing zebrafish at 5 fish per liter [13]. However, there are many institutions that consider a higher density, up to 20 fish per liter, as an acceptable housing density [14,15,16]. One study determined a zebrafish housing density of up to 12 fish per liter had no negative impact on reproductive performance [17]. Primary factors determining an appropriate housing density include food access, water quality, pathogens, and mating opportunities; all of which would not play a role in very brief housing for euthanasia [18]. With that said, there can still be aggression between conspecifics that were not housed in the same tanks and animals should not be euthanized with fish from separate tanks. While we saw no behavioral differences in the fish euthanized at a density of 5 fish/L compared with 10 fish/L in Eto-6, the times to LORR (median times; 17 sec *vs* 24 sec) and undulation cessation (median times; 30 sec *vs* 60.5 sec) were significantly shorter in fish euthanized at 5 fish/L, respectively. This aligned with our hypothesis that fish euthanized at a lower density would have a shorter euthanasia time, potentially resulting from higher etomidate concentrations being absorbed as less fish are absorbing the agent. However, we saw no significant differences in the times to COO (median times; 106 sec and 112 sec) and loss of startle reflex (median times; 540 sec and 720 sec), respectively. A limiting factor was the small sample size (n=5 *vs* n=10) and further research may elucidate whether density plays a largely significant role. Regardless of densities, all fish were euthanized effectively at 30 min.

Etomidate provides a viable alternative to euthanize adult zebrafish because it is simple to use, stable, and non-toxic to humans. The most common anesthetic agent overdose used for euthanasia, MS-222, has several drawbacks. MS-222 is a powder requiring a fume-hood for preparation due to potential human health hazards. These hazards include skin, eye, and respiratory irritation and long-term occupational exposure is associated with retinal toxicity. To minimize these health hazards, MS-222 must be prepared while wearing proper PPE such as gloves and goggles. MS-222 also requires the addition of bicarbonate to increase the pH, since it can be very acidic and, therefore, irritating to the fish. Even when properly prepared many researchers report their fish experiencing aversion to MS-222 [4,19]. Etomidate would only require a syringe and needle to dispense, minimizing not only human safety risks but also medical waste in the form of PPE. Etomidate is also a pharmaceutical grade compound, shelf stable for 24 months, and is not light sensitive, compared to MS-222, that should not be exposed to light and loss of potency occurs between three and 10 days of storage at room temperature [20,21]. While etomidate has a long shelf life and is easy to store, one of the drawbacks could potentially be the price. In the United States a bottle of etomidate can cost roughly 32$ for 100 ml [22]. However, using a 10 mg/L concentration this equates to about 30 cents for 1 ml and you would only be paying roughly 1.50$ per liter. Etomidate is also commonly used in human medical facilities in vials that are commonly discarded after single use. Animal facilities that work closely with human hospitals may be able to use the excess vials for zebrafish euthanasia and avoid any costs whatsoever. Another factor to consider is etomidate’s ease of availability, due to various manufacturing companies responsible for production. There is a single FDA approved manufacturer of MS-222, Syndel USA, Ferndale WA (Tricaine-S), which may cause availability issues [23].

Another anesthetic agent with a similar mechanism of action to etomidate, propofol, results in a quick LORR and COO but due to the agent’s opaqueness there can be difficulty assessing potentially fine aversive behaviors such as increased operculation [24,25]. Another recent paper investigated metomidate hydrochloride, a rapid acting nonbarbiturate that is also related to etomidate. Their findings paralleled ours with fish losing the righting reflex within one minute and losing operculation within a couple minutes. Similarly, they found fish responded to vibration, sound, or touch for upwards of 66 minutes; similar to the increased time to loss of the startle reflex we observed. However, they assessed ornamental fish and used significantly higher concentrations of metomidate [26]. Another paper had used conditioned aversion testing and found metomidate to be less aversive than MS-222 [19]. Metomidate is considered an acceptable method of euthanasia in Canada and Europe [27,28]. Currently metomidate hydrochloride is not listed in the AVMA euthanasia guidelines as an acceptable euthanasia method due its status as an FDA-indexed drug [2]. Presently, etomidate is available and may provide a viable alternative in the United States.

One variable that we did not consider in this study was the time the fish were exposed to etomidate. Amend et al. evaluated the use of etomidate as an anesthetic agent and found 0% mortality in adult zebrafish immersed at a 3.0 mg/L concentration exposed for 2 minutes but a 20% mortality when exposed for 20 minutes at the same 3.0 mg/L concentration. They also found that adult zebrafish immersed at 15 mg/L had a 95% mortality when exposed for just 2 minutes [29]. This variation emphasizes the need to evaluate a variety of concentrations in combination with a variety of exposure times in finding the quickest and least aversive euthanasia method using an anesthetic overdose.

## Conclusions

This study evaluated two separate etomidate concentrations and two separate fish densities to euthanize adult zebrafish. Fish euthanized in Eto-10 had shorter times to undulation cessation, COO, and loss of startle reflex than fish euthanized in Eto-6. Fish euthanized at 5 fish/L had shorter times to LORR and undulation cessation than fish euthanized at 10 fish/L. We conclude either Eto-6 or Eto-10 is efficacious at euthanizing zebrafish when immersed for at least 30 min, but an adjunct method of euthanasia should still be performed to confirm brain death, although no zebrafish recovered from the etomidate exposure at either etomidate concentration. We also conclude that fish may be euthanized at 5 or 10 fish/L, but 5 fish/L may provide quicker immobilization times. Further studies may address alternative concentrations, immersion times, densities, lifestages, as well as histological impact on tissues. This data provides more information on an alternative anesthetic agent overdose as an adult zebrafish euthanasia method providing quick immobilization without signs of aversive behavior.

## Acknowledgments

We would like to thank the Stanford Department of Comparative Medicine for supporting this research project.

